# Mathematical Modeling of PCR with Variable Concentrations of DNA and *Taq* Polymerase

**DOI:** 10.1101/2023.03.16.532908

**Authors:** Fraz Ahmad, Saeeda Zia, Rashid Saif, Muhammad Hassan Raza

## Abstract

Polymerase Chain Reaction (PCR) is one of the important techniques in molecular biology for many diagnostics and research applications. In this current theoretical PCR optimization research, out of its seven recipe reagents, two of the important constituents e.g., DNA which might be very rare in forensic cases and polymerase enzyme due to its high cost were mathematically modeled and computationally simulated using MATLAB programming platform. Chemical equations of PCR stages were established using the law of mass action followed by differential equations derivations from each of the DNA amplification phases. Firstly, a mathematical model of *Taq* polymerase enzyme with its variable concentrations ranging from 0.05-0.35 *IU*/*μL* was formulated which showed the best PCR product of 0.61 *ng/μL* obtained with 0.2 *IU/μL* of polymerase. Similarly, by fixing the optimal polymerase concentration obtained in the previous simulation, the prime-DNA variable concentrations ranging from 0.5-3.5 *ng/μL* were further simulated which showed the best PCR product of 0.94 *ng/μL* at 2.0 *ng/μL* of DNA. PCR wet-lab experimentation and gel-electrophoresis visualization validated mathematically simulated optimized concentrations of the aforementioned two reagents which vary the PCR product considerably. Current findings indicate that mathematically optimized concentrations of *Taq*-polymerase and template-DNA may reduce the PCR cost, time and energy, but other factors of target-sequence specificity, purity/concentrations of the other reagents, working hygiene, technical intricacies, biosafety and biosecurity factors may also contribute the desired yield of the amplicon.

## 1 Introduction

The invention and development of polymerase chain reaction (PCR) has revolutionized the experimental science in incredible ways [1]. PCR uses enzymatic activity of polymerase to make billions of copies from targeted DNA region in short period of time. Prior to this discovery, the scientists were limited to analyze DNA due to restricted sample availability. PCR has a broad impact across the field and has led researchers to gain deeper insights in understanding processes in molecular biology, evolutionary biology, medical diagnostics and forensic analysis [2].

In 1985, Kary Mullis invented the PCR technique which consist of three temperature-induced steps including denaturation, annealing and extension [3]. These three steps can be repeated from 25-40 cycles depending on the required yield. In denaturation step, the sample containing double-stranded DNA is separated into two single strands by breaking hydrogen bonds at 94°C. The short single stranded DNA primer is then allowed to bind to the targeted region on DNA at the annealing stage. The temperature for this stage varies from 50-65°C depending on the GC content of the primers and other factors. Primers are specific in nature and only bind to the region for which they have been designed. In third stage the thermostable *Taq_pol_* form new DNA strand by adding deoxyribonucleotide (dNTPs) at 3’-end of primer in Primer-DNA complex. This enzyme catalyzes the formation of growing strand by forming a phosphodiester bond. The optimal temperature of thermostable *Taq_pol_* driven extension phase is 72°C [4].

In past PCR has been mathematically modelled using variety of approaches. Jim et.al developed a mathematical model using the law of mass action and simplified the assumptions regarding the structure of reactions using differential and chemical equations. They concluded that dynamically optimizing the extension and annealing stages of individual samples may significantly reduce the total time for a PCR run [5]. Another research in similar domain investigated the thermal effect on reaction behavior and optimized the temperature for primer binding by using Michaelis–Menten equation [6]. A study proposed a mathematical model of PCR kinetics curve, they simulated the PCR process over a series of reaction cycles which allows for predicting the efficiency of real-time PCR at any cycle number [7]. In all these studies, researchers predicted model for growth per cycle, reaction time, temperature effect on PCR, kinetic and its efficiency. However, there is no mathematical model available for optimizing PCR reagents to our information.

The aim of current research is to develop a mathematical model for PCR with variable concentrations of DNA and *Taq_pol_* while considering other reagents constant. These two reagents are important to model as *Taq_pol_* is commercially expensive, and secondly there may be a shortage of DNA concentration in certain forensic samples [8]. The concentration of *Taq_pol_* and template DNA was mathematically modelled using law of mass action and simulation using MATLAB. The law of mass action was used to drive the chemical equations which further supported the derivation of differential equations. After solving the differential equations analytically, MATLAB’s ‘ode45’ package was used to run simulations to obtain optimized concentrations of amplicon, along with graphical representations. Finally, these predictions were validated through wet-lab PCR amplification experiments. Our wet and dry lab simulations might be helpful for determining the optimal concentrations of the aforementioned reagents which will provide other researchers the opportunity to obtain optimal PCR product with minimal ultrapure DNA and *Taq_pol_* concentration.

## 2 PCR dynamics

### 2.1 Denaturation

Double stranded DNA is typically held together by hydrogen bonds which are formed between nitrogenous bases of opposite strands. Denaturation is caused by thermal induced dissociation of double-stranded DNA (dsDNA) into single stranded DNA (ssDNA). A sample containing dsDNA is subjected to a high temperature e.g., (94°C) which breaks the hydrogen bonds between DNA resulting into two single strands multiple times during the reaction. The chemical equation of this reversible DNA denaturation can be written as.

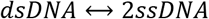

This equation represents the conversion of *dsDNA* into two *ssDNA* molecules multiple times.

### 2.2 Annealing

The specific primers are designed to anneal to the single-stranded DNA (ssDNA) templates at their complementary region when allowed to react at their respective temperature. A primer is a short complementary sequence when it binds to ssDNA it forms the primer-DNA complex also known as primed-ssDNA. The annealing step chemical reaction equation can be written as.

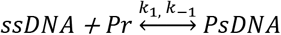

Where *k*_1_ and *k*_–1_ is the rate of forward and reverse reaction respectively. *Pr* is the primer while *PsDNA* is the primed-ssDNA complex. In actual PCR dynamics scenario, the optimum temperature is applied to maximize the forward reaction as compared to reverse reaction.

### 2.3 Extension

The *Taq_pol_* extracted from *Thermus aquaticus* bacteria binds to primed-ssDNA at 3’-end to perform the extension by adding nucleotides sequentially and form dsDNA at 72°C. The overall chemical reaction of extension can be written as.

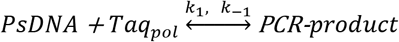

The *k*_1_ and *k*_–1_ is the rate of forward and reverse reaction. The above “*PCR-product*” variable is denoted with *“P”* in the downstream equations.

The equation for individual step in adding single nucleotide to growing chain is shown below where *“C”* represents the complex of primed-ssDNA and *Taq_pol_,* the *“B”* represents the nucleotide base and *C1, C2, C3* … *Cn* represents the *primed-ssDNA-Taq_pol_* complex with added nucleotides.

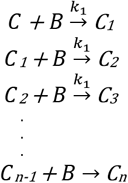

*Taq_pol_* enzyme make it easier to add bases in the correct order from the sense strand 5’ first nucleotide of forward primer while on the antisense strand from 5’ first nucleotide of reverse primer as shown in the annealing step of Fig. 1.

**Fig. 1.**
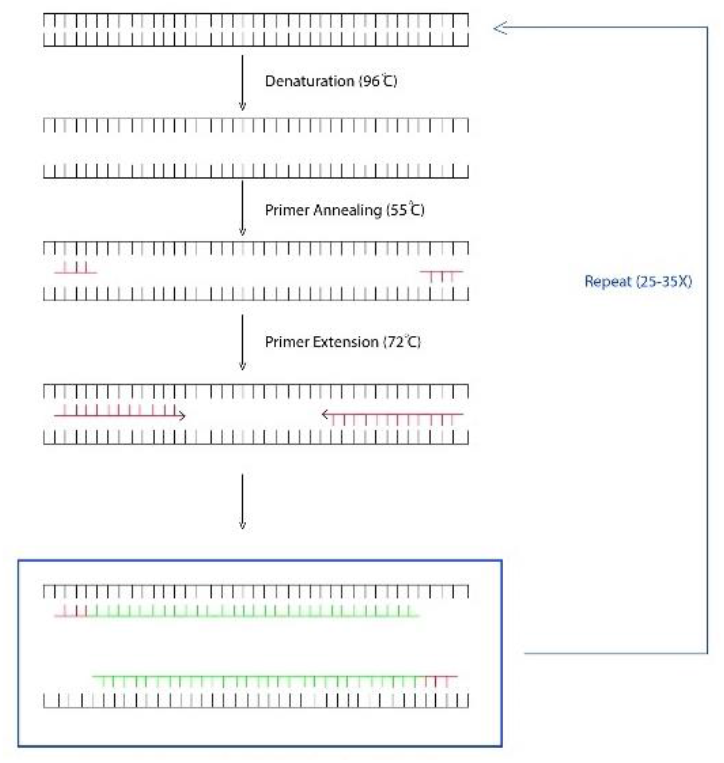
PCR amplification is represented by denaturation followed by annealing and extension steps.

## 3 Model formulation of *Taq_pol_*

PCR relies on seven basic reagents to amplify DNA with *Taq_pol_* being the key reagent that facilitates the synthesis of new strands. Modeling the quantity of *Taq_pol_* is essential for gaining deeper insights into reaction kinetics and achieving maximum efficiency, as well as predicting cost-effective concentrations to improve the PCR protocol. The volume and concentration of *Taq_pol_* which were used in the construction of following model is as follows and mentioned in the Table 1.

**Table 1.**
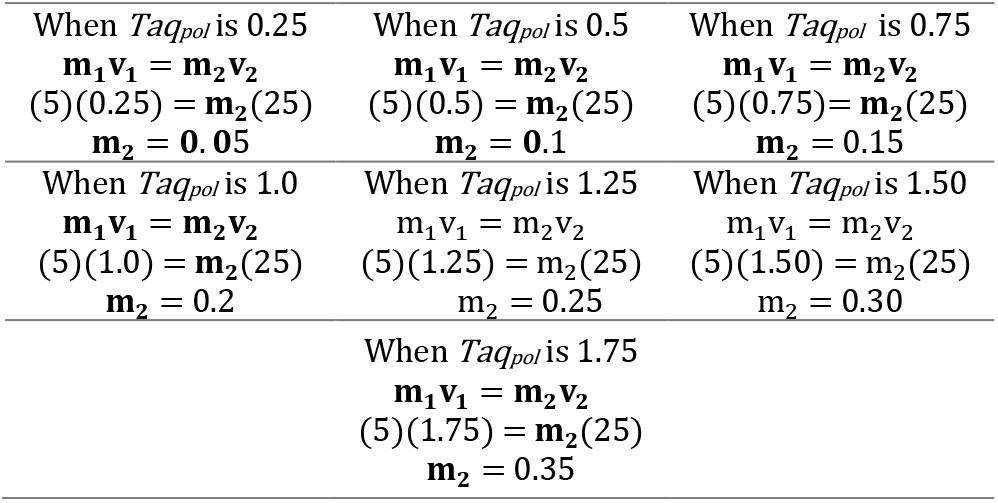
Calculation of *Taq_pol_* final concentration

The attributes used for modeling *Taq_pol_* concentration is given in Table 2. 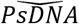 represent the initial amount of primed-ssDNA. *k*_1_ and *k_–1_* are the rate constants for the forward and reverse reactions respectively. These estimates are based on published data or experimental results obtained through the PCR protocol [5]. Variable concentration of *Taq_pol_* is taken for simulation and represented as 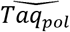.

**Table 2.** Variable symbols, concentrations and units used in the model construction for *Taq_pol_*

| Symbol | Concentration | Units |
| --- | --- | --- |
| $\overline{PsDNA}$ | 0.7 | ng/μL |
| $\overline{Taq_{pol}}$ | 0.05, 0.1, 0.15, 0.20, 0.25, 0.30, 0.35 | IU/μL |
| $k_1$ | 0.205 | μL/μmol/sec |
| $k_{-1}$ | 0.01025 | μL/μmol/sec |

### 3.1 Annealing

Law-of-mass-action is used to construct the differential equations of annealing stage.

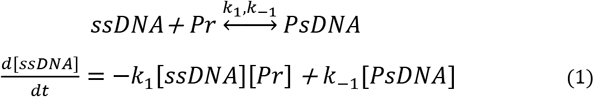

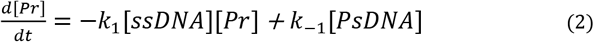

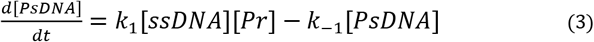

The *k*_1_ is the rate of forward reaction and *k*_–1_ is the rate of reverse reaction. *“Pr”* is the concentration of primer, *“ssDNA”* is the concentration of single strand DNA and *“PsDNA”* is the concentration of primed-ssDNA. Eq. (1), (2) and (3) represents the change in the concentrations of ssDNA, primer and primed-ssDNA with respect to time.

### 3.2 Extension

The law-of-mass-action is used to derive differential equations that characterize the variation in concentration of each reactant and products.

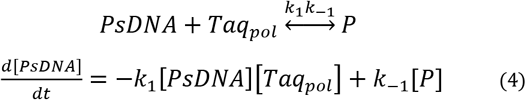

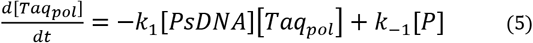

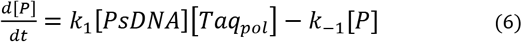

The “*P*” is concentration of product and “*Taq_pol_*” is the concentration of *Taq_pol_*, “*PsDNA*” Is the concentration of primed-ssDNA. Eq. (1), (2) and (3) represents the change in the concentrations of primed-ssDNA, *Taq_pol_* and product. Our equations can be simplified by law of conservation.

By adding Eq. (4) and Eq. (6)

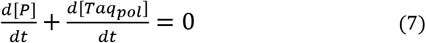

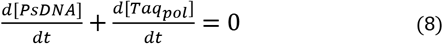

This gives rise two constant quantities of *M_1_* and *M_2_* in the below equations.

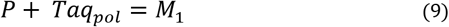

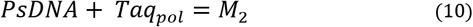

By using above formulations, the system can be reduced to a single equation.

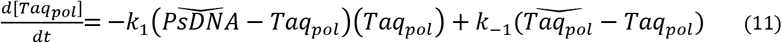

The initial conditions are as follows,

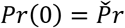 (the concentration of primer at the beginning of the reaction)
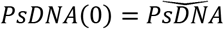 (the concentration of primed-ssDNA at the beginning of the reaction)
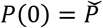 (the concentration of product at the beginning of the reaction)

After solving Eq. (11) by separation of variables. The solution is,

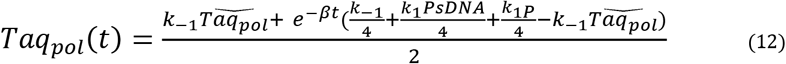

Where,

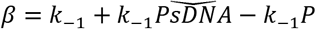

The different concentrations of *Taq_pol_* are used to get optimized value for maximum yield to make PCR more efficient. The first cycle starts with primers, *Taq_pol_*, DNA to be copied along with other reagents and their values should be in accordance to values used in the wet-lab experiments. The final equation (12) provides the information about the concentration of *Taq_pol_* and product over time during PCR. Where *Taq_pol_*(*t*) represents the concentration of *Taq_pol_* at time t, 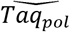 represents the initial concentration of *Taq_pol_*, *k*_–1_ is the rate of reverse reaction, *PsDNA* represents the concentration of primed-ssDNA, P represents the concentration of product and β is a constant.

## 4 Model formulation of primed-ssDNA

The amount of primed-ssDNA in a PCR has a significant effect on the PCR outcome. Sometime the amount of primed-ssDNA is not ideal for required procedure, such as in crime scene investigations where the amount of DNA may not sufficient for analysis. Therefore, it is important to mathematically model primed-ssDNA to achieve maximum amplicon. The primed-ssDNA variable concentrations are modelled by fixing optimal polymerase concentration obtained from the previous simulation. The concentration of *PsDNA* which were used in the construction of following model is as follows and mentioned in the Table 3.

**Table 3.**
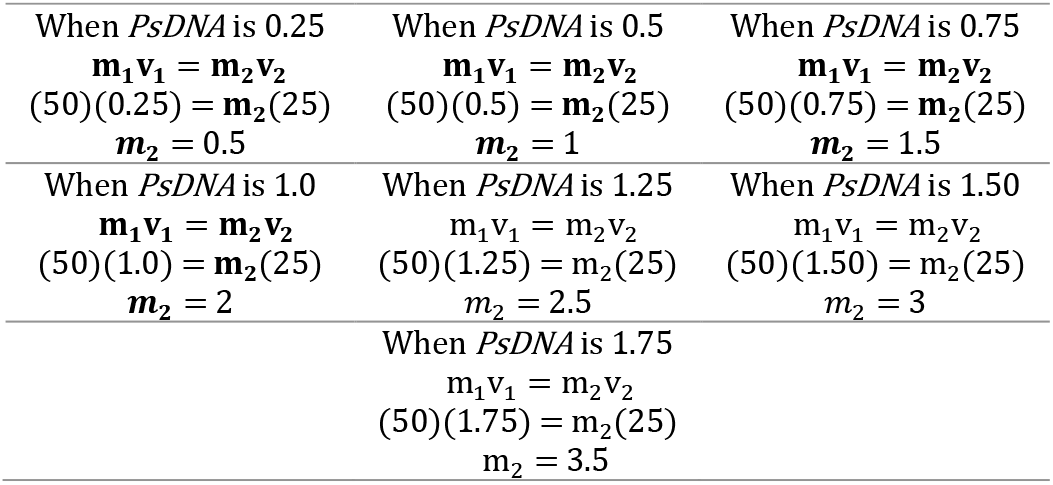
Calculation of *PsDNA* final concentrations

The variables attributes used for modeling *PsDNA* concentration are provided in Table 4. Where 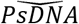 represent the initial amount of primed-ssDNA. *k1* and *k-1* are the rate constants for the forward and reverse reactions respectively, which were obtained from the literature [5]. Variable concentration of *PsDNA* is taken for simulation and represented as 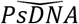.

**Table 4.** Variable symbols, their concentrations and unit used in the model construction for *PsDNA*

| Symbol | Concentrations | Units |
| --- | --- | --- |
| $\overline{Taq_{pol}}$ | 1.0 | IU/μL |
| $\overline{PsDNA}$ | 0.5, 1.0, 1.5, 2.0, 2.5, 3.0, 3.5 | ng/μL |
| $k_1$ | 0.205 | μL/μmol/sec |
| $k_{-1}$ | 0.01025 | μL/μmol/sec |

For primed-ssDNA modeling the differential equations is written with the help of law of mass action.

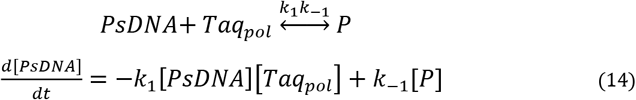

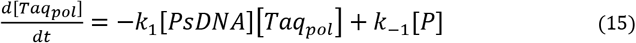

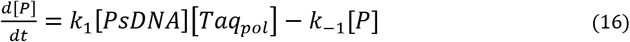

The *k*_1_ is the rate of forward reaction and *k*_–1_ is the rate of reverse reaction. “P” is the concentration of product and 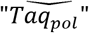 is the concentration of *Taq_pol_*, 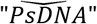 Is the concentration of primed-ssDNA. Eq. (14), Eq. (15) and Eq. (16) represents the change in the concentrations of primed-ssDNA, *Taq_pol_* and product.

The equations can be simplified by conservation law

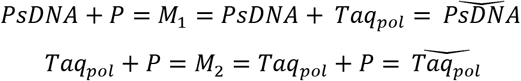

The *M*_1_ and *M*_2_ are constant. Using above equations, the system can be simplified to a single equation.

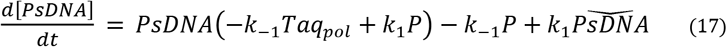

The initial conditions are,

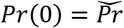 (the concentration of primer at the beginning of the reaction)
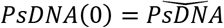 (the concentration of primed-ssDNA at the beginning of the reaction)
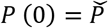 (the concentration of product at the beginning of the reaction)

After solving equation by separation of variables. The solution is

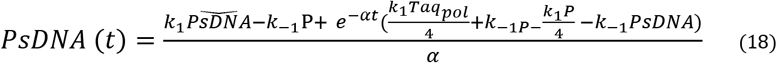

Where,

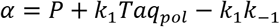

Different quantities of initial primed-ssDNA were used to predict the optimal concentration. The reaction was initiated with the addition of primers and *Taq_pol_*, followed by DNA to be amplified and other reagents. The concentrations of these reagents were adjusted based on the values used in the experiments. The final equation (18) provides information about concentration of primed-ssDNA and product concentration over time during PCR. Where *PsDNA* (t) represents the concentration of primed-ssDNA at time t, 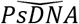 represents the initial concentration of primed-ssDNA, k_1_ is the rate of forward reaction, *Taq_pol_* represents the concentration of *Taq_pol_*, P represents the concentration of product and *α* is a constant.

## 5 MATLAB simulation for *Taq_pol_*

The *Taq_pol_* quantities ranging 0.05-0.35 *IU/μL* were simulated using MATLAB which suggested that maximum PCR product of 0.6690 *ng*/μ*L* may be achieved using 0.35 *IU*/μ*L* of *Taq_pol_*, but 0.2 *IU*/μ*L* concentration of *Taq_pol_* was found satisfactory and optimal by producing 0.6116 *ng*/μ*L* PCR product due to unproportionate increase of the product as compared to the increase in the quantity of *Taq_pol_*. MATLAB package ode45 was used for theoretical simulation of product. The concentrations of product obtained using variable concentration of *Taq_pol_* are given in Table 5, where shaded row shows the optimal suggested concentrations of *Taq_pol_*.

**Table 5.** Theoretical estimations of variable concentration of *Taq_pol_* and PCR product

| Conc. of $Taq_{pol}$<br>(IU/μL) | Vol. of $Taq_{pol}$<br>(μL) | Vol. of PsDNA<br>(μL) | Conc. of Product<br>(ng/μL) |
| --- | --- | --- | --- |
| 0.05 | 0.25 | 0.7 | 0.2230 |
| 0.1 | 0.50 | 0.7 | 0.4117 |
| 0.15 | 0.75 | 0.7 | 0.5416 |
| 0.20 | 1.0 | 0.7 | 0.6116 |
| 0.25 | 1.25 | 0.7 | 0.6443 |
| 0.30 | 1.50 | 0.7 | 0.6602 |
| 0.35 | 1.75 | 0.7 | 0.6690 |

### 5.1 Graphical representation of *Taq_pol_* simulation

A mathematical simulation of above reagent depicted through graphical representation which shows dynamic behavior of the PCR product with initial fixed concentration of 0.7*μL* PsDNA and variable concentrations of *Taq_pol_* as with PCR progression (Figure 2). The best considered PCR product is shown in Figure 2d where product is 0.6116 *ng/μL* with 0.2 *IU/μL of Taq_pol_*.

**Fig. 2.**
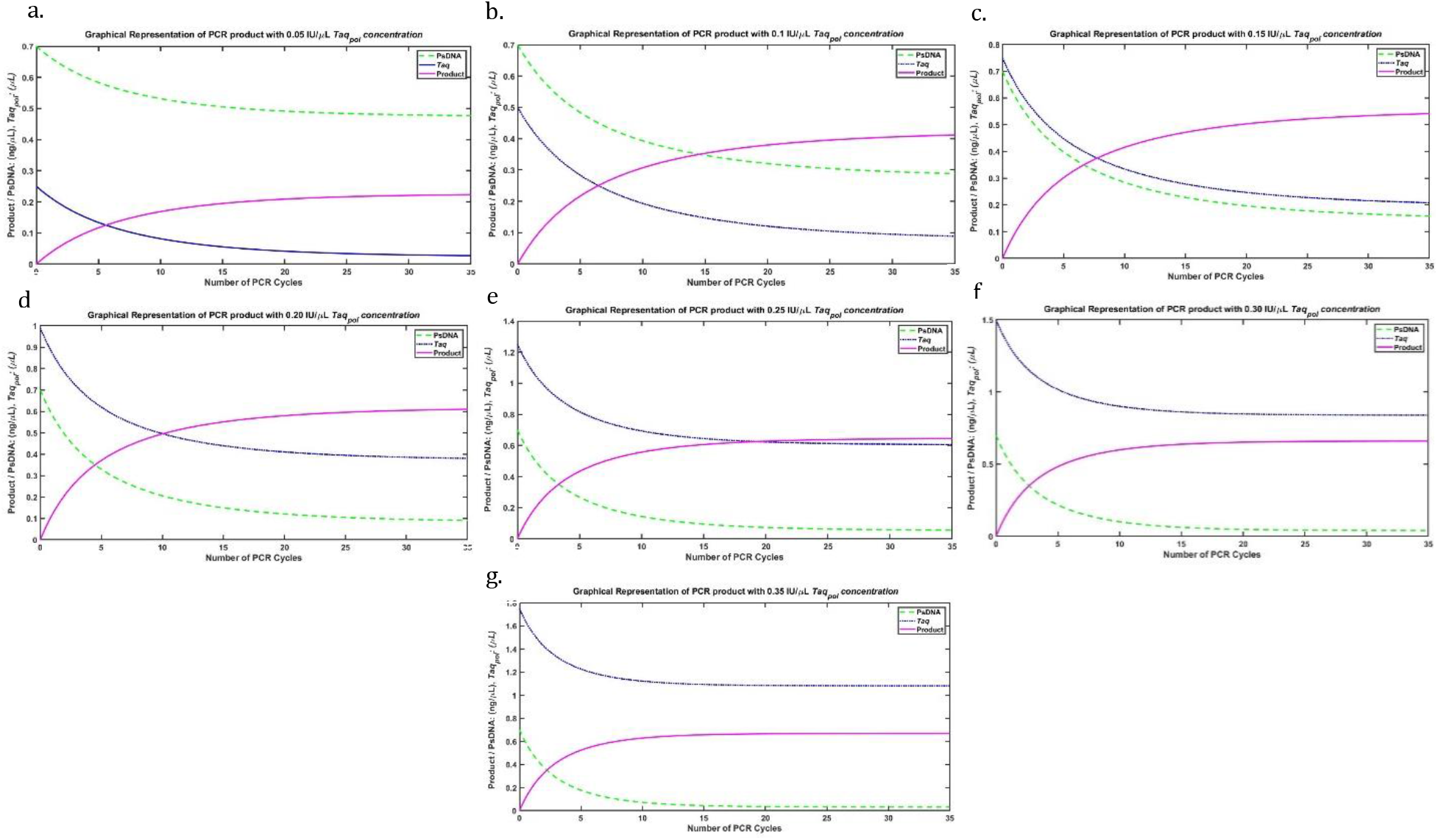
Graphical representation of predicted product concentration with variable *Taq_pol_*, in which *Taq_pol_*, PCR product and *PsDNA* are represented with blue, green and purple lines respectively. **a)** The concentration of *PsDNA* and unreactive *Taq_pol_* decreases causing the increase PCR product from 0 to 0.2230 *ng/μL* **b)** The concentration of *PsDNA* and unreactive *Taq_pol_* decreases causing the increase in PCR product from 0 to 0.4117 *ng/μL* **c)** The concentration of *PsDNA* and unreactive *Taq_pol_* decreases causing the increase in PCR product from 0 to 0.5416*ng/μL* **d)** The concentration of *PsDNA* and unreactive *Taq_pol_* decreases causing the increase in PCR product from 0 to 0.6116 *ng/μL* **e)** The concentration of *PsDNA* and unreactive *Taq_pol_* decreases causing the increase in PCR product from 0 to 0.6443 *ng/μL* **f)** The concentration of *PsDNA* and unreactive *Taq_pol_* decreases causing the increase PCR product from 0 to 0.6602*ng/μL* **g)** Similarly, the concentration of *PsDNA* and unreactive *Taq_pol_* decreases causing the increase of PCR product from 0 to 0.6690*ng/μL.*

### 5.2 Wet-lab validation of simulated PCR with *Taq_pol_*

The predicted concentration obtained from the mathematical simulation was validated through wet-lab PCR experiment. Genomic DNA was extracted and specific primers were designed against the target sequence. Seven PCR reactions were conducted on single sample using same concentration of PCR reagents within each tube except different concentration of *Taq_pol_* which were previously simulated as shown in the Table 5. Gel electrophoresis was used to have an idea of the product concentration by comparing band strength of PCR amplicon. The wet-lab results confirmed the optimal product was obtained at 0.2 *IU/μL* of *Taq_pol_* (boxed band in Figure 4) as predicted mathematically. Although the last three wells have sharp bands but using proportionately higher concentration of *Taq_pol_* which has slight impact on the product concentration while first three band has low product.

**Fig. 4.**
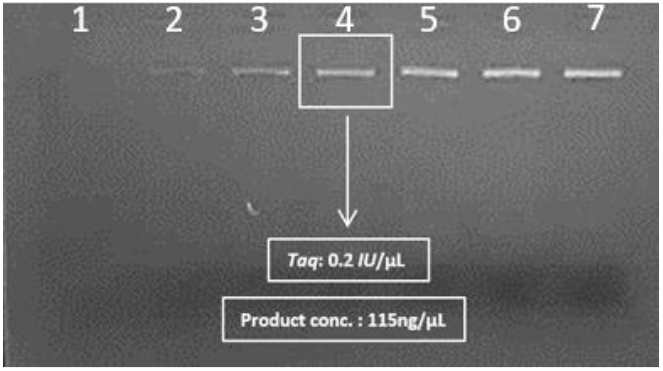
PCR run using variable *Taq_pol_* concentrations from 0.05 to 0.35 /U/μL

## 6 MATLAB simulation for primed-ssDNA

The primed-ssDNA quantities ranging 0.5-3.5 *ng/μL* were simulated for maximum PCR yield using previously optimized 0.2 *IU/μL* concentration of *Taq_pol_*. The model suggested the maximum PCR product of 0.9989*ng/μL* may be achieved using 3.5*ng/μL PsDNA,* but 2.0*ng*/μ*L* concentration of *PsDNA* was found satisfactory and optimal by producing 0.9490*ng*/μ*L* PCR product due to unproportionate increase of the product as compared to the increase in the quantity of *PsDNA.* MATLAB package ode45 was used for theoretical simulation of product. The concentrations of product obtained using variable concentration of *PsDNA* are given in Table 6, where shaded row in the table shows the best considered concentrations of *PsDNA.*

**Table 6.**
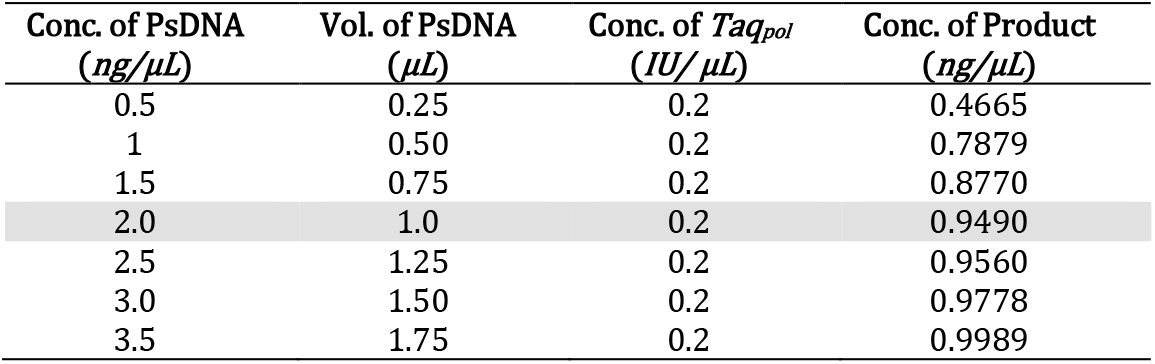
Theoretical estimations of variable concentration of *PsDNA* and PCR product

| Conc. of $PsDNA$<br>($ng/\mu L$) | Vol. of $PsDNA$<br>($\mu L$) | Conc. of $Taq_{pol}$<br>($IU/\mu L$) | Conc. of Product<br>($ng/\mu L$) |
| --- | --- | --- | --- |
| 0.5 | 0.25 | 0.2 | 0.4665 |
| 1 | 0.50 | 0.2 | 0.7879 |
| 1.5 | 0.75 | 0.2 | 0.8770 |
| 2.0 | 1.0 | 0.2 | 0.9490 |
| 2.5 | 1.25 | 0.2 | 0.9560 |
| 3.0 | 1.50 | 0.2 | 0.9778 |
| 3.5 | 1.75 | 0.2 | 0.9989 |

### 6.1 Graphical representation of *PsDNA* simulation

A mathematical simulation of above reagent depicted through graphical representation which shows dynamic behavior of the PCR product with initial fixed concentration of 0.2*IU/μL Taq_pol_* and variable concentrations of *PsDNA* as with PCR progression (Figure 3). The best considered PCR product is shown in Figure 3d where product is 0.9490*ng/μL* with 2.0*ng/μL of Taq_pol_*.

**Fig. 3.**
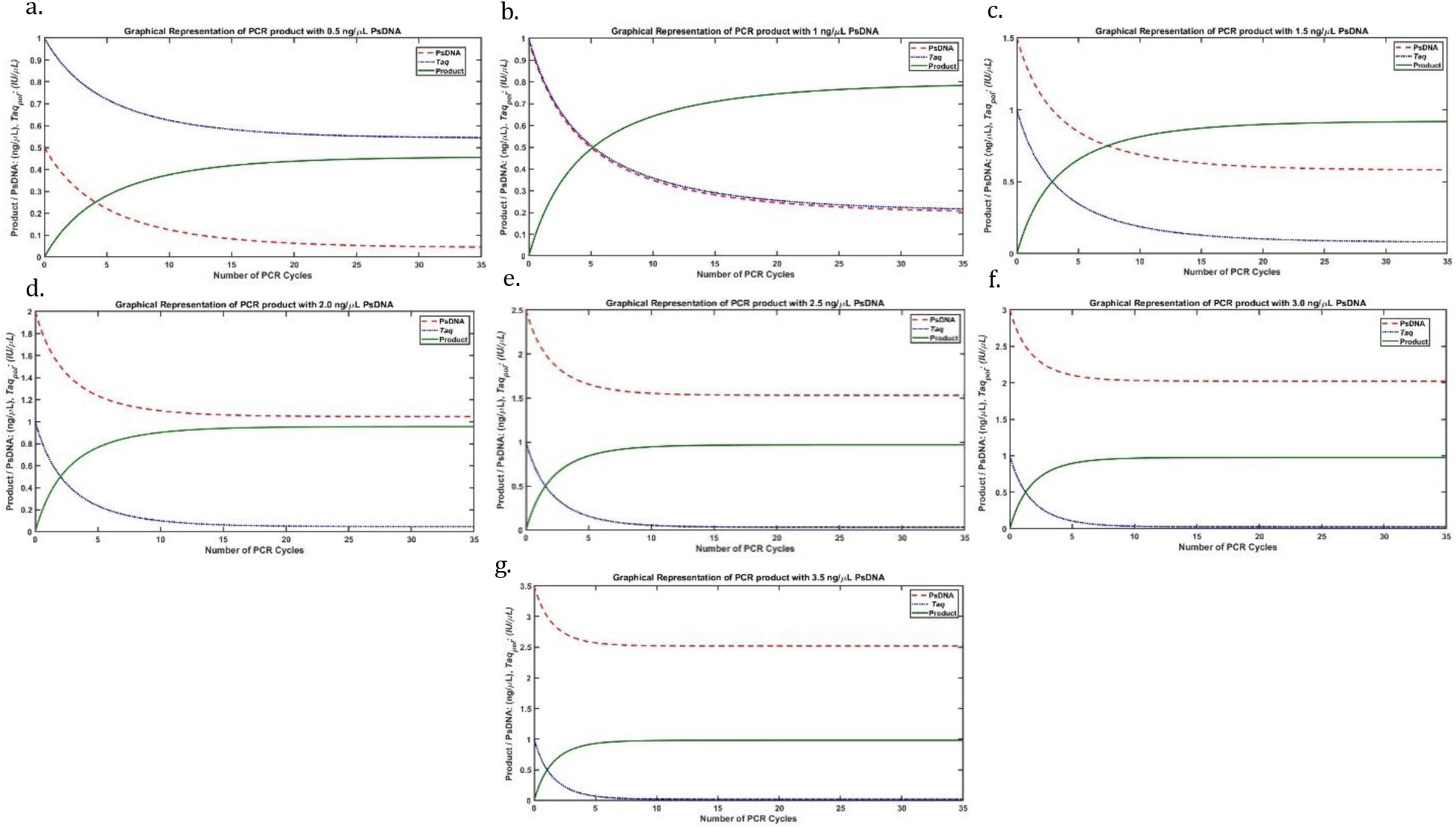
Graphical representation of predicted product concentration with variable *PsDNA,* in which *Taq_pol_, PsDNA* and PCR product are represented with purple, orange and green lines respectively. **a)** The concentration of *PsDNA* and unreactive *Taq_pol_* decreases causing the increase in PCR product from 0 to 0.4665*ng/μL* **b)** The concentration of *PsDNA* and unreactive *Taq_pol_* decreases causing the increase in PCR product from 0 to 0.7879*ng/μL* **c)** The concentration of *PsDNA* and unreactive *Taq_pol_* decreases causing the increase in PCR product from 0 to 0.8770*ng/μL* **d)** The concentration of *PsDNA* and unreactive *Taq_pol_* decreases causing the increase in PCR product from 0 to 0.9490*ng/μL* **e)** The concentration of *PsDNA* and unreactive *Taq_pol_* decreases causing the increase in PCR product from 0 to 0.9560*ng/μL* **f)** The concentration of *PsDNA* and unreactive *Taq_pol_* decreases causing the increase in PCR product from 0 to 0.9778*ng/μL* **g)** Similarly, the concentration of *PsDNA* and unreactive *Taq_pol_* decreases causing the increase in PCR product from 0 to 0.9989*ng/μL.*

### 6.2 Wet-lab validation of simulated PCR with primed-ssDNA

The predicted concentration obtained from the mathematical simulation was validated through wet-lab experiments. Single sample genomic DNA was used to perform seven PCR reactions that were conducted using 0.2*IU/μL* fixed concentration of *Taq_pol_* with different concentration of *PsDNA* which were previously simulated as shown in the Table 6. Gel electrophoresis was run to have an idea of the product concentration by comparing band strength of PCR amplicon. The wet-lab results confirmed the optimal product was obtained at 2.0*ng/μL* of *PsDNA* (boxed band in Figure 5) as predicted mathematically. Although the last three wells have sharp bands but using proportionately higher concentration of *PsDNA* which has slight impact on the product concentration while first three band has negligible product.

**Fig. 5.**
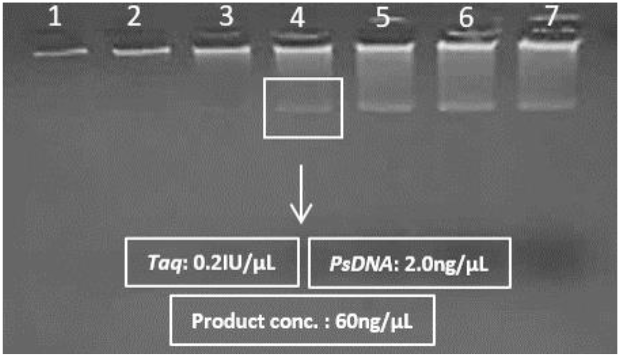
PCR run using *PsDNA* concentrations from 0.5 to 3.5 *ng/μL*

## 7 Discussion

PCR has become one of the most extensively used procedures in molecular biology. The number of PCR based applications has grown significantly, impacting cancer genetics and pathogen identification [9]. In this research, we determined the optimal conditions for PCR that can be achieved through mathematical modeling, resulting in maximum PCR yield with minimum *Taq_pol_* and template *PsDNA*. The high cost and limited availability of PCR reagents have prompted the need for a cost-effective and efficient approach to optimize the reaction conditions. Mathematically modeling of the reaction making it possible to reduce the concentrations of reagents during the wet-lab experiments, thereby making the overall process more cost-effective and optimized with minimal DNA concentration possible.

Our findings indicates that maximum PCR product can be obtained when reaction is performed with *Taq_pol_* and primed-ssDNA at concentration of 0.2 *IU/μL* and 2.0 *ng/μL* respectively by assuming other reagents as constant. The results obtained from our wet-lab experiments were found to be consistent with the predictions made by mathematical model, indicating a high degree of conformity between the two. This phenomenon highlights the effectiveness of the mathematical approach in accurately describing the reaction dynamics of the PCR and importance of incorporating mathematical modeling in the molecular biology studies.

*Taq_pol_* and *PsDNA* concentrations higher than optimized value results only in slight increase in PCR product. Using an excessive *Taq_pol_* may not significantly increase the yield of PCR product. This occurs when an excessive amount of *Taq_pol_* competes for the same template DNA, leading to the formation of inefficient complexes that block access of the polymerase to the DNA. Moreover, the 5’-3’ exonuclease activity of the enzyme may also result in the formation of smear artifacts [10]. In the scenario where too little template is used, the reaction may not be sensitive enough to detect the target sequence and yield a sufficient amount of product. Alternatively, if too much template is used, the reaction may become saturated leading to reduced yields or even inhibiting the amplification. This is because a high concentration of template DNA can compete with the primers for binding sites, making it more difficult for the primers to bind and initiate the amplification process [11].

Similar mathematical prediction studies on PCR efficiency have been done in past by optimizing factors like temperature, reaction kinetics and multiplex assays [12]. However, there is no such model available that can predict the optimal concentration of DNA and *Taq_pol_* for maximum PCR efficiency. But one of another modelling approach was used to understand the underlying processes and factors involved in annealing and extension kinetics. The results of this study suggested that dynamically optimizing the annealing and extension stages of individual samples may significantly reduce the total time for a PCR run [13].

There are few limitations of this study, as the PCR optimization is based on two variables while assuming concentration of other reagents fixed. Predicting a model with low number of variables can make it simpler and easier to interpret, but it can also lead to lower accuracy. This can overcome by predicting a model with high number of variables but it will make model more complex and inclined to partial differential equation (PDE) problem. This highlights the need for further endeavors to develop a more comprehensive mathematical model that takes into account the effects of all the relevant variables in the PCR reaction.

## 8 Conclusion

This study suggests that optimal PCR product may be achieved with 0.2 *IU/μL Taq_pol_* and 2.0 *ng/μL PsDNA* of ultrapure quality, but at the same time concentrations of other reagents may interfere the PCR product, so a further model may be proposed by considering other reagent concentrations in action rather than assuming as constants, which will enable researchers to make informed decisions about the efficacy and cost-effectiveness of PCR.

